# Identification of pathogenic missense mutations using protein stability predictors

**DOI:** 10.1101/2020.06.11.146068

**Authors:** Lukas Gerasimavicius, Xin Liu, Joseph A Marsh

**Affiliations:** MRC Human Genetics Unit, Institute of Genetics and Molecular Medicine, University of Edinburgh, Edinburgh EH4 2XU, UK

## Abstract

Attempts at using protein structures to identify disease-causing mutations have been dominated by the idea that most pathogenic mutations are disruptive at a structural level. Therefore, computational stability predictors, which assess whether a mutation is likely to be stabilising or destabilising to protein structure, have been commonly used when evaluating new candidate disease variants, despite not having been developed specifically for this purpose. We therefore tested 12 different stability predictors for their ability to discriminate between pathogenic and putatively benign missense variants. We find that one method, FoldX, considerably outperforms all others in the identification of disease variants. Moreover, we demonstrate that employing absolute energy change scores improves performance of nearly all predictors. Importantly, however, we observe that the utility of computational stability predictors is highly heterogeneous across different proteins, and that they are all are inferior to the best performing variant effect predictors for identifying pathogenic mutations. We suggest that this is largely due to alternate molecular mechanisms other than protein destabilisation underlying many pathogenic mutations. Thus, better ways of incorporating protein structural information and molecular mechanisms into computational variant effect predictors will be required for improved disease variant prioritisation.

## Introduction

Advances in next generation sequencing technologies have revolutionised research of genetic variation, increasing our ability to explore the basis of human disorders and enabling huge databases covering both pathogenic and putatively benign variants^1,2^. Novel sequencing methodologies allow the rapid identification of variation in the clinic and are helping facilitate a paradigm shift towards precision medicine^3,4^. Despite this, however, it remains challenging to distinguish the small fraction of variants with medically relevant effects from the huge background of mostly benign human genetic variation.

A particularly important research focus is single nucleotide variants that lead to amino acid substitutions at the protein level, *i.e*. missense mutations, which are associated with more than half of all known inherited diseases^5,6^. Thus, a large number of computational methods have been developed for the identification of potentially pathogenic missense mutations, *i.e*. variant effect predictors. Although different approaches vary in their implementation, a few types of information are most commonly used, including evolutionary conservation, changes in physiochemical properties of amino acids, biological function, known disease association and protein structure^7^. While these predictors are clearly useful for variant prioritisation, and show a statistically significant ability to distinguish known pathogenic from benign variants, they still make many incorrect predictions^8–10^, and the extent to which we can rely on them for diagnosis remains limited^11^.

An alternative approach to understanding the effects of missense mutations is with computational stability predictors. These are programs that have been developed to assess folding or protein-protein interaction energy changes upon mutation (change in Gibbs free energy – ΔΔG in short). This can be achieved by approximating structural energy through linear physics-based pairwise energy scoring functions, their empirical and knowledge-based derivatives, or a mixture of such energy terms. Statistical and machine learning methods are employed to parametrise the scoring models. These predictors have largely been evaluated against their ability to predict experimentally determined ΔΔG values. Great effort has been previously made to assess stability predictor performance in producing accurate or well-correlated energy change estimates upon mutation, as well as assessing their shortfalls, such as biases arising from destabilising variant overrepresentation in training sets and lack of self-consistency predicting forward-backward substitutions^12–18^. Moreover, their practical utility has been demonstrated through their extensive usage in the fields of protein engineering and design^19–21^.

Although computational stability predictors have not been specifically designed to identify pathogenic mutations, they are very commonly used when assessing candidate disease mutations. For example, publications reporting novel variants will often include the output of stability predictors as evidence in support of pathogenicity^22–25^. This relies essentially upon the assumption that the molecular mechanism underlying many or most pathogenic mutations is directly related to the structural destabilisation of protein folding or interactions^26–29^. However, despite their widespread application to putatively pathogenic variants, there has been little to no systematic assessment of computational stability predictors for their ability to predict disease mutations. A number of studies have assessed the real-world utility for individual protein targets and families using certain stability predictors^30–34^. However, numerous computational stability predictors have now been developed and, overall, we still do not have a good idea of which methods perform best for the identification of disease mutations, and how they compare relative to other computational variant effect predictors.

In this work, we explore the applicability and performance of 12 methodologically diverse structure-based protein stability predictors for distinguishing between pathogenic and putatively benign missense mutations. We find that FoldX substantially outperforms other stability predictors for the identification of disease mutations, and also to demonstrate the value of using absolute ΔΔG values to account for potentially stabilising mutations. However, this work also highlights the limitations of stability predictors for predicting disease, as they still miss many pathogenic mutations and perform worse than many variant effect predictors, thus emphasising the likely importance of considering alternate molecular disease mechanisms beyond protein destabilisation.

## Results

We tested 12 different computational stability predictors on the basis of accessibility, automation or batching potential, computation speed, as well as recognition – and included FoldX^35^, Rosetta^35^, PoPMusic^36^, I-Mutant^37^, SDM^38^, SDM2^39^, mCSM^40^, DUET^41^, CUPSAT^42^, MAESTRO^43^, ENCoM^44^ and DynaMut^45^ (**Table 1**). We ran each predictor against 13,508 missense mutations from 96 different high-resolution (< 2 Å) crystal structures of monomeric proteins. This included 3,338 missense variants from ClinVar^2^ annotated as pathogenic or likely pathogenic, and 10,170 variants observed in the human population, taken from gnomAD^1^. Each protein structure had at least 10 pathogenic missense mutations that could be modelled with the stability predictors. While it is possible that some of the gnomAD variants could be damaging under certain circumstances (*e.g*. if observed in a homozygous state, if they cause late-onset disease, or there is incomplete penetrance), the large majority of them are likely to be non-pathogenic, and we therefore refer to them as “putatively benign”.

**Table 1.**
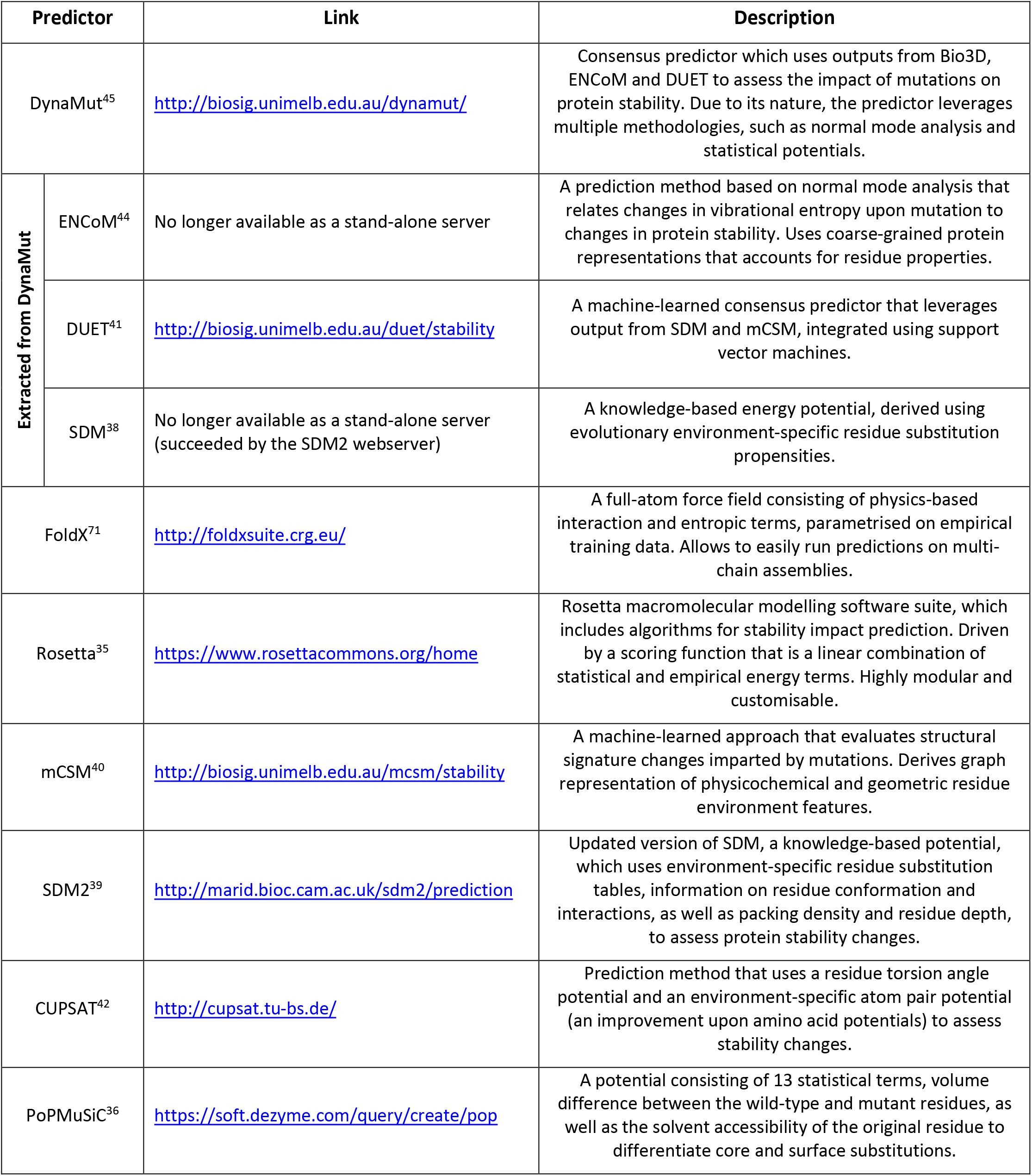
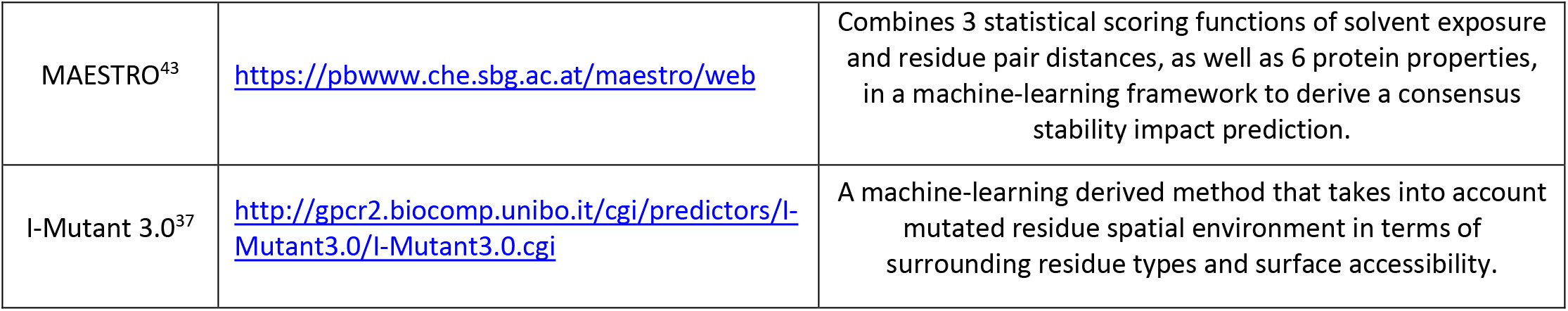
Protein stability predictors used in this study.

| Predictor |  | Link | Description |
| --- | --- | --- | --- |
| DynaMut<sup>45</sup> |  | http://biosig.unimelb.edu.au/dynamut/ | Consensus predictor which uses outputs from Bio3D, ENCoM and DUET to assess the impact of mutations on protein stability. Due to its nature, the predictor leverages multiple methodologies, such as normal mode analysis and statistical potentials. |
| Extracted from DynaMut | ENCoM<sup>44</sup> | No longer available as a stand-alone server | A prediction method based on normal mode analysis that relates changes in vibrational entropy upon mutation to changes in protein stability. Uses coarse-grained protein representations that accounts for residue properties. |
|  | DUET<sup>41</sup> | http://biosig.unimelb.edu.au/duet/stability | A machine-learned consensus predictor that leverages output from SDM and mCSM, integrated using support vector machines. |
|  | SDM<sup>38</sup> | No longer available as a stand-alone server (succeeded by the SDM2 webserver) | A knowledge-based energy potential, derived using evolutionary environment-specific residue substitution propensities. |
| FoldX<sup>71</sup> |  | http://foldxsuite.crg.eu/ | A full-atom force field consisting of physics-based interaction and entropic terms, parametrised on empirical training data. Allows to easily run predictions on multi-chain assemblies. |
| Rosetta<sup>35</sup> |  | https://www.rosettacommons.org/home | Rosetta macromolecular modelling software suite, which includes algorithms for stability impact prediction. Driven by a scoring function that is a linear combination of statistical and empirical energy terms. Highly modular and customisable. |
| mCSM<sup>40</sup> |  | http://biosig.unimelb.edu.au/mcsm/stability | A machine-learned approach that evaluates structural signature changes imparted by mutations. Derives graph representation of physicochemical and geometric residue environment features. |
| SDM2<sup>39</sup> |  | http://marid.bioc.cam.ac.uk/sdm2/prediction | Updated version of SDM, a knowledge-based potential, which uses environment-specific residue substitution tables, information on residue conformation and interactions, as well as packing density and residue depth, to assess protein stability changes. |
| CUPSAT<sup>42</sup> |  | http://cupsat.tu-bs.de/ | Prediction method that uses a residue torsion angle potential and an environment-specific atom pair potential (an improvement upon amino acid potentials) to assess stability changes. |
| PoPMuSiC<sup>36</sup> |  | https://soft.dezyme.com/query/create/pop | A potential consisting of 13 statistical terms, volume difference between the wild-type and mutant residues, as well as the solvent accessibility of the original residue to differentiate core and surface substitutions. |

|  |  |  |
| --- | --- | --- |
| MAESTRO<sup>43</sup> | https://pbwww.che.sbg.ac.at/maestro/web | Combines 3 statistical scoring functions of solvent exposure and residue pair distances, as well as 6 protein properties, in a machine-learning framework to derive a consensus stability impact prediction. |
| I-Mutant 3.0<sup>37</sup> | http://gpcr2.biocomp.unibo.it/cgi/predictors/I-Mutant3.0/I-Mutant3.0.cgi | A machine-learning derived method that takes into account mutated residue spatial environment in terms of surrounding residue types and surface accessibility. |

To investigate the utility of the computational stability predictors for the identification of pathogenic missense mutations, we used receiver operating characteristic (ROC) plots to assess the ability of ΔΔG values to distinguish between pathogenic and putatively bening mutations (**Fig. 1A**). This was quantifed by the area under the curve (AUC), which is equal to the probability of a randomly chosen disease mutation being assigned a higher-ranking score than a random benign one. Of the 12 tested structurebased ΔΔG predictors, FoldX clearly performs the best as a predictor of human missense mutation pathogenicity, with an AUC value of 0.661. The next best methods, Rosetta and PoPMuSiC, show considerably worse, with ranking probabilities of 0.617 and 0.614. Evaluating the performance differences through bootstrapping we found that Rosetta and PoPMuSiC significantly underperform compared to FoldX, with p-values of 1 x 10^-7^ and 8 x 10^-9^, respectively, while the difference between them was insignificant. The remaining predictors demonstrated a wide range of lower performance values.

**Figure 1:**
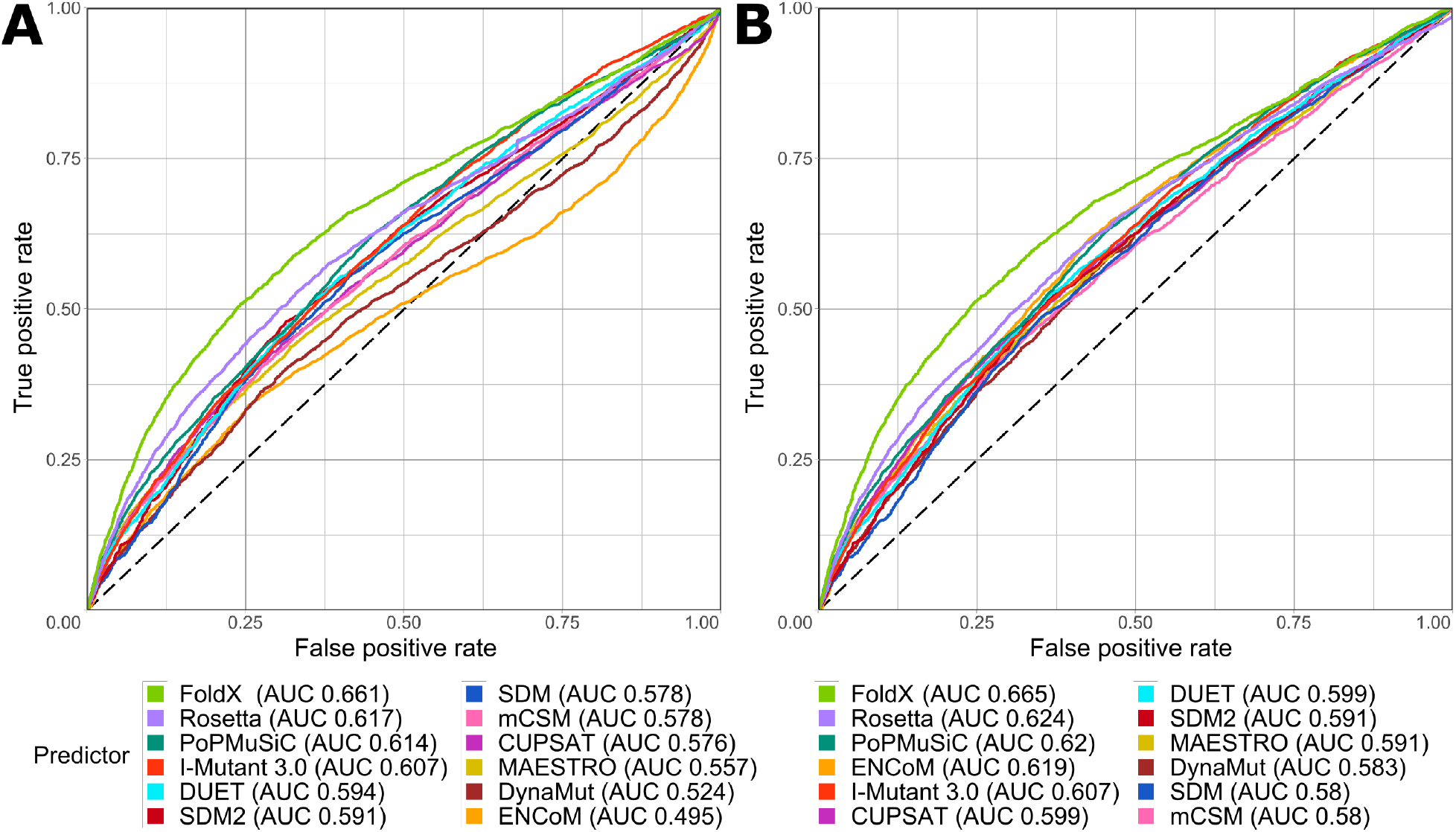
Using ΔΔG values from protein stability predictors to discriminate between pathogenic and putatively benign missense variants. Receiver operating characteristic (ROC) curves are plotted for each predictor, with the classification performance being presented next to its name in the form of area under the curve (AUC). **A)** ROC curves for classification performance using native ΔΔG value scale for each predictor. **B)** ROC curves for predictor classification performance when using absolute ΔΔG values.

Two predictors, ENCoM and DynaMut, stand out for their unusual pattern in the ROC plots, with a rotated sigmoidal shape where the false positive rate becomes greater than the true positive rate at higher levels. Close inspection of the underlying data shows that this is indicative of the predicted energy change distribution tails for the disease-associated class extending both directions away from the putatively benign missense mutation score density. This suggests that a considerable portion of pathogenic missense mutations are predicted by these predictors to excessively stabilise the protein.

While the analysis **Fig. 1A** assumes that protein destabilisation should be indicative of mutation pathogenicity, it also possible for stabilising mutations to be pathogenic^46,47^. Therefore, we repeated the analysis using absolute ΔΔG values, so that both destabilising and stabilising mutations are treated equivalently (**Fig. 1B**). This improved the performance of most predictors, while not reducing the performance of any. The most drastic change was observed for ENCoM, which improved from worst to fourth best predictor, with an increase in AUC from 0.495 to 0.619. However, the top three predictors, FoldX, Rosetta and PoPMuSiC, improve only slightly and do not change in ranking.

Using the ROC point distance to the top-left corner^48^, we establish the best disease classification ΔΔG value for each predictor when assessing general perturbation (**Table 2)**. It is interesting to note that FoldX demonstrates the best classification performance when utilising 1.58 kcal/mol as the stability change threshold, which is remarkably to the value of 1.5 kcal/mol previously suggested and used in a number of other works when assessing missense mutation impact on stability^13,33,49^. Of course, these threshold values should be considered far from absolute rules, and there are many pathogenic and benign mutations above and below the thresholds for all predictors. For example, nearly 40% of pathogenic missense mutations have FoldX values lower than the threshold, whereas approximately 35% of putatively benign variants are above the threshold.

**Table 2.** Best stability predictor classification thresholds according to ‘distance-to-corner’ metric. The performance metrics and their 95% confidence intervals were derived from 2000 bootstraps of the data.

| Predictor | Absolute $\Delta\Delta G$ threshold | False positive rate (95% confidence interval) | True positive rate (95% confidence interval) |
| --- | --- | --- | --- |
| FoldX | 1.578 | 0.339–0.357 | 0.591–0.623 |
| Rosetta | 1.886 | 0.390–0.409 | 0.572–0.605 |
| PoPMuSiC | 0.795 | 0.417–0.437 | 0.584–0.618 |
| CUPSAT | 1.455 | 0.415–0.434 | 0.549–0.583 |
| MAESTRO | 0.321 | 0.418–0.437 | 0.544–0.578 |
| SDM | 1.025 | 0.350–0.370 | 0.477–0.511 |
| SDM2 | 0.875 | 0.365–0.385 | 0.510–0.544 |
| mCSM | 0.889 | 0.433–0.453 | 0.542–0.575 |
| DUET | 0.803 | 0.400–0.421 | 0.548–0.582 |
| I-Mutant 3.0 | 0.915 | 0.405–0.424 | 0.545–0.578 |
| ENCoM | 0.221 | 0.415–0.436 | 0.598–0.632 |
| DynaMut | 0.476 | 0.446–0.467 | 0.570–0.605 |

We also calculated AUC values for each protein separately and compared the distribution across predictors (**Fig. 2**). FoldX again performs much better than other stability predictors for the identification of pathogenic mutations, with a mean ROC of 0.681, compared to Rosetta at 0.627, PoPMuSiC at 0.621, and ENCoM at 0.630. Interestingly, the protein-specific performance was observed to be extremely heterogeneous across all predictors. While some predictors performed extremely well (AUC > 0.9) for certain proteins, each predictor has a considerable number of proteins for which they perform worse than random classification (AUC < 0.5).

**Figure 2:**
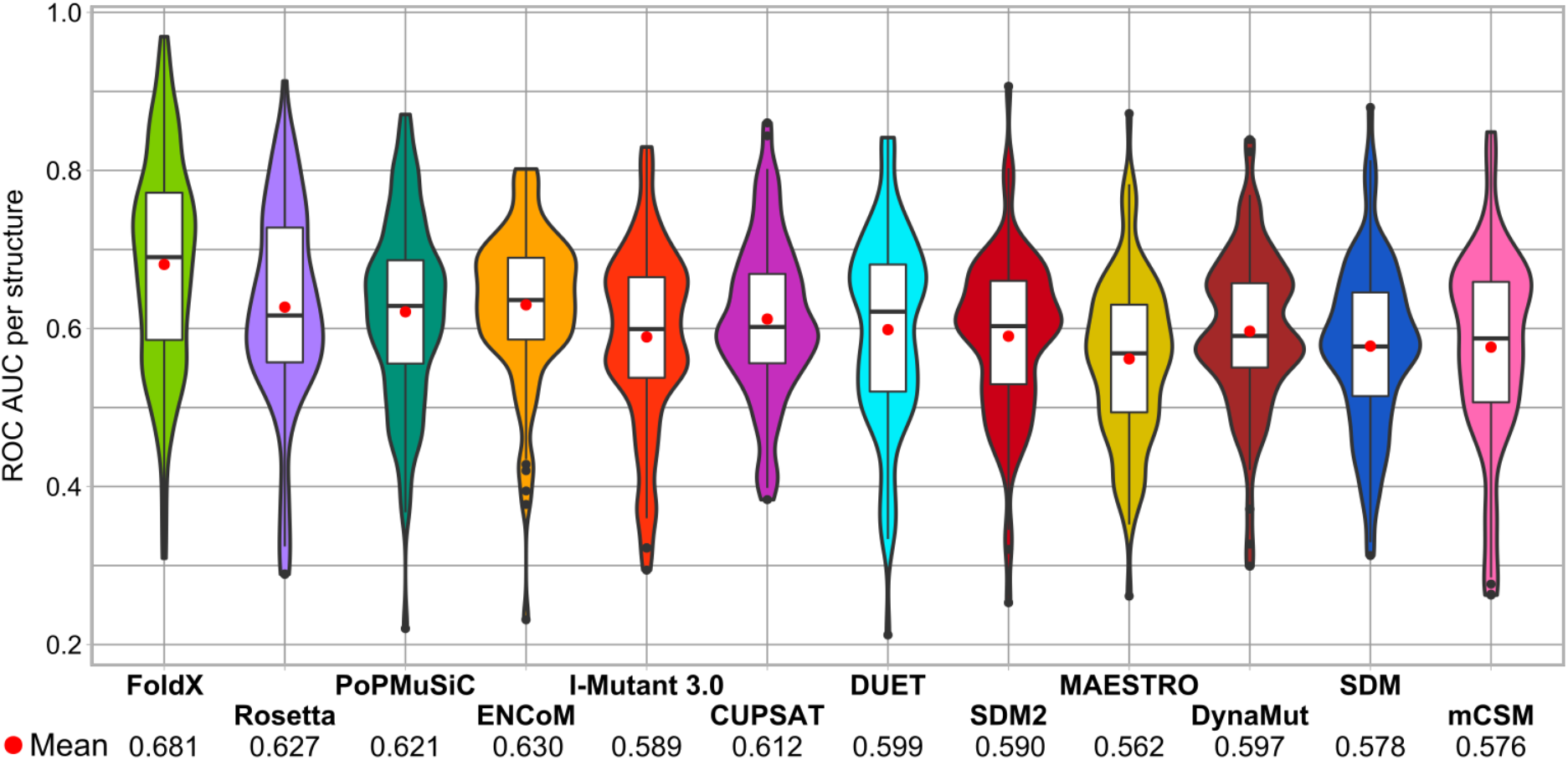
The heterogeneity of protein-specific missense variant classification performance. All the stability predictors exhibit very high degrees of heterogeneity in their protein-specific performance, as measured by the ROC AUC on a per-protein basis. Absolute ΔΔG values were used during protein-specific tool assessment. The mean performance of each predictor is indicated by a red dot and numerically showcased below the plot. Boxes inside the violins illustrate the interquartile range (IQR) of the protein-specific performance points, with the whiskers measuring 1.5 IQR. Boxplot outliers are designated by black dots.

Finally, we compared the performance of protein stability predictors to a variety of different computational variant effect predictors **(Fig. 3)**. Importantly, we excluded any predictors trained using supervised learning techniques, as well as meta-predictors that utilise the outputs of other predictors, thus including only predictors we labelled as unsupervised and empirical in our recent study^10^. This is due to the fact that predictors based upon supervised learning are likely to have been directly trained on some of the same mutations used in our evaluation dataset, making a fair comparison impossible^10,50^. A few predictors perform substantially better than FoldX, with the best performance seen for SIFT4G^51^, a modified version of the SIFT algorithm^52^. Interestingly, FoldX is the only stability predictor to outperform the BLOSUM62 substitution matrix^53^. On the other hand, all stability predictors performed better than a number of simple evolutionary constraint metrics.

**Figure 3:**
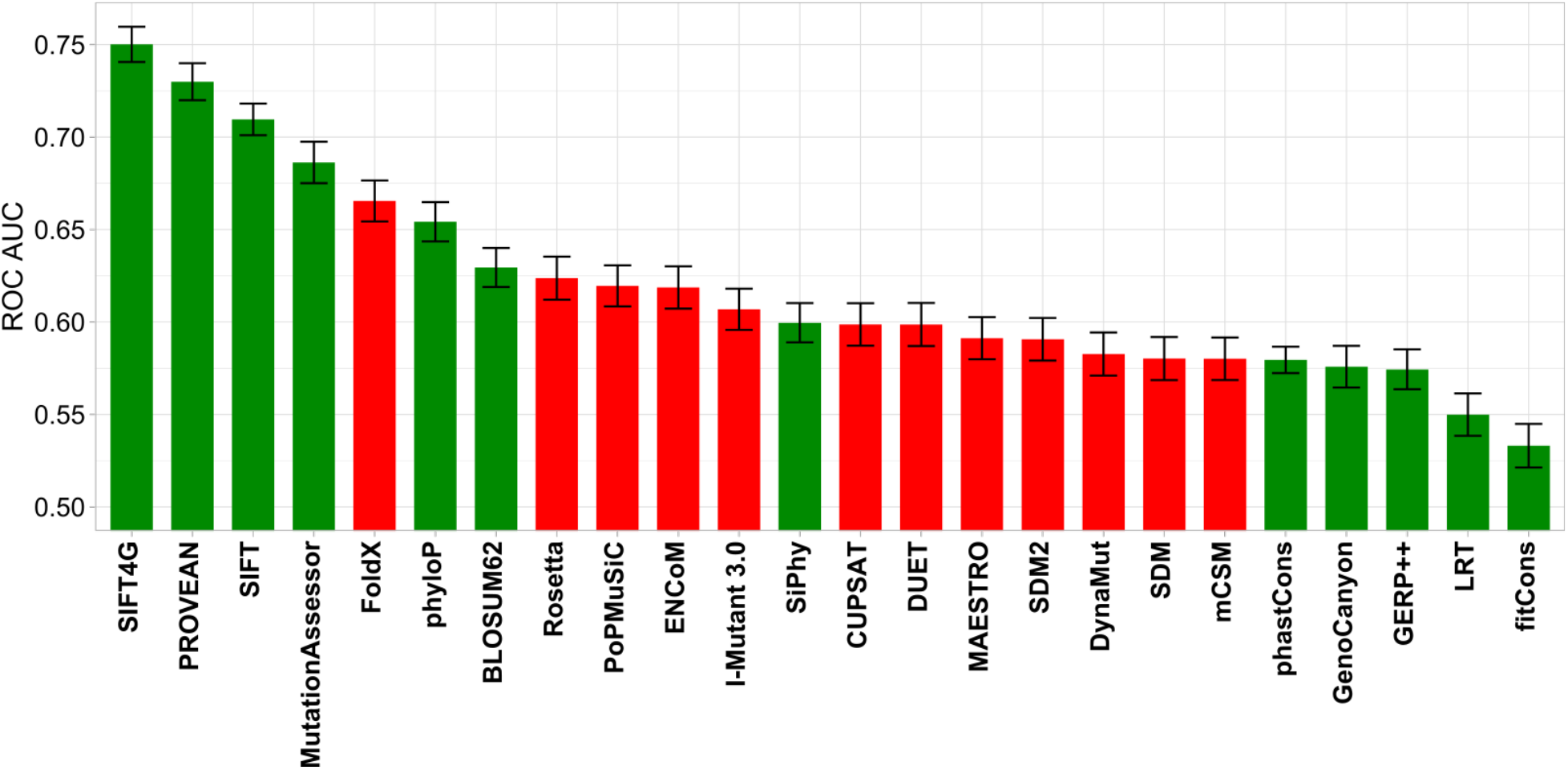
Performance comparison of protein stability and variant effect predictors for identifying pathogenic variants. Error bars indicate the 95% confidence interval of the ROC AUC as derived through bootstrapping. Stability predictors are shown in red, while other variant effect prediction methods are shown in green. Absolute ΔΔG values were used for stability-based methods.

## Discussion

The first purpose of this study was to compare different computational stability predictors for their ability to identify pathogenic missense mutations. In this regard, FoldX is the winner, clearly outperforming the other ΔΔG prediction tools. It also has the advantage of being computationally undemanding, fairly easy to run, and flexible in its utilisation. Compared to other methods that employ physics-based terms, FoldX introduces a few unique energy terms into its potential, notably the theoretically derived entropy costs for fixing backbone and side chain positions^54^. However, the main reason behind its success is likely the parametrisation of the scoring function, resulting from the well optimised design of the training and validation mutant sets, which aimed to cover all possible residue structural environments^55^. Interestingly, while the form of the FoldX function, consisting of mostly physics-based energy terms, has not seen much change over the years, newer knowledge-based methods, which leverage statistics derived from the abundant sequence and structure information, demonstrate poorer and highly varied performance. However, it is important to note that the performance of FoldX does not necessarily mean that it is the best predictor of experimental ΔΔG values or true destabilisation, as that is not what we are testing here.

There are two factors likely to be contributing to the improvement in the identification of pathogenic mutations using absolute ΔΔG values. First, while most focus in the past has been on destabilising mutations, some pathogenic missense mutations are known to stabilise protein structure. As an example, the H101Q variant of chloride intracellular channel 2 (CLIC2) protein, which is thought to play a role in calcium ion signalling, leads to developmental disabilities, increased risk to epilepsy and heart failure^56^. The CLIC2 protein is soluble, but requires insertion into the membrane for its function, with a flexible loop connecting its domains being functionally implicated in a necessary conformational rearrangement. The histidine to glutamine substitution, which occurs in the flexible loop, was predicted to have an overall stabilising energetic effect due to conservation of weak hydrogen bonding, but also the removal of charge that the protonated histidine exerted on the structure^56^. The ΔΔG predictions were followed up by molecular dynamics simulations, which supported the previous conclusions by showing reduced flexibility and movement of the N-terminus, with functional assays also revealing reduced membrane integration of the CLIC2 protein in line with the rigidification hypothesis^57^. However, other interesting examples of negative effects of over-stabilisation exist in enzymes and protein complexes, manifesting through the activity-stability trade-off, rigidification of co-operative subunit movements, dysregulation of protein-protein interactions, and turnover^46,47,58^.

In addition, it may be that some predictors are not as good at predicting the direction of the change in stability upon mutation. That is, they can predict structural perturbations that will be reflected in the magnitude of the ΔΔG value, but are less accurate in their prediction of whether this will be stabilising or destabilisng. For example, ENCoM and DynaMut predict nearly half of pathogenic missense mutations to be stabilising (41% and 44%, respectively), whereas FoldX predicts only 13%. While FoldX, Rosetta and PoPMuSiC are all driven by scoring functions consisting of a linear combination of physics- and statistics-based energy terms, ENCoM is based on normal mode analysis, and relates the assessed entropy changes around equilibrium upon mutation to the state of free energy. DynaMut, a consensus method, integrates the output from ENCoM and several other predictors (**Table 1**) into its score^45^. The creators of ENCoM found that their method is less biased at predicting stabilising mutations^59^. From our analysis, we are unable to say anything about what proportion of pathogenic mutations are stabilising *vs* destabilising, or about which methods are better at predicting the direction of stability change, but this is clearly an issue that needs more attention in the future.

The second purpose of our study was to try to understand how useful protein stability predictors are for the identification of pathogenic missense mutations. Here, the answer is less clear. While all methods show some utility for discriminating between pathogenic and benign variants, it is notable and perhaps surprising that all methods except FoldX performed worse than the simple BLOSUM62 substitution matrix, which suggests that these methods may be relatively limited utility for variant prioritisation. Even FoldX was unequivocally inferior to multiple variant effect predictors, suggesting that it should not be relied upon by itself for the identification of disease mutations.

One reason for the limited success of stability predictors in the identification of disease mutations is that predictions of ΔΔG values are still far from perfect. For example, a number of studies have compared ΔΔG predictors, showing heterogeneous correlations with experimental values on the order of R=0.5 for many predictors^12,13,60^. However, a recent work has also revealed problems with the noise in experimental stability data used to benchmark the prediction methods, generally assessed through correlation values^61^. Taking noise and data distribution limitations into account, it is estimated that with currently available experimental data the best ΔΔG predictor output correlations should be in the range 0.7-0.8, while higher values would suggest overfitting^61^. As such, even assuming that ‘true’ ΔΔG values were perfectly correlated with mutation pathogenicity, we would still expect these computational predictors to misclassify many variants.

The existence of alternate molecular mechanisms underlying pathogenic missense mutations is also likely to be a major contributor to the underperformance of stability predictors compared to other variant effect predictors. At the simplest level, our analysis does not consider intermolecular interactions. Thus, given that pathogenic mutations are known to often occur at protein interfaces and disrupt interactions^62,63^, the stability predictors would not be likely to identify these mutations in this study. We tried to minimise the effects of this by only considering crystal structures of monomeric proteins, but the existence of a monomeric crystal structure does not mean that a protein does not participate in interactions. Fortunately, FoldX can be easily applied to protein complex structures, so the effects of mutations on complex stability can be assessed.

Pathogenic mutations that act via other mechanisms are also likely to be missed by stability predictors. For example, we have previously shown that dominant-negative mutations in ITPR1^64^ and gain-of-function mutations in PAX6^65^ tend to be mild at a protein structural level. This is consistent with the simple fact that highly destabilising mutations would not be compatible with dominant-negative or gain-of-function mechanisms. Similarly, hypomorphic mutations that cause only a partial loss of function are also likely to be less disruptive to protein structure than complete loss-of-function missense mutations^66^.

These varying molecular mechanisms are all likely to be related to the large heterogeneity in predictions we observe for different proteins in Fig 2. Similarly, the specific molecular and cellular contexts of different proteins could also limit the utility of ΔΔG values for predicting disease mutation. For example, even weak perturbations in haploinsufficient proteins could lead to a deleterious phenotype. At the same time, intrinsically stable proteins, proteins that are overabundant or functionally redundant could tolerate perturbing variants without such high ΔΔG variants being associated with disease. Finally, in some cases, mildly destabilising mutations can unfold local regions, leading to proteasome mediated degradation of the whole protein^32,34,67^.

There could be considerable room for improvement in ΔΔG predictors and their applicability to disease mutation identification. Recently emerged hybrid methods, such as VIPUR^68^ and SNPMuSiC^69^, show promise of moving in the right direction, as they assess protein stability changes upon mutation while attempting to increase the interpretability and accuracy by taking the molecular and cellular contexts into account. However, none of the mentioned hybrid methods employ FoldX, which, given our findings here, may be a good strategy. Rosetta is also promising due to its tremendous benefit demonstrated in protein design. It should be noted that the protocol used for Rosetta in our work utilised rigid backbone parameters, due to the computation costs and time constraints involved in allowing backbone flexibility. An accuracy-oriented Rosetta protocol, or the “cartesian_ddg” application in the Rosetta suite, which allows structure energy minimisation in Cartesian space, may lead to better performance^35,70^.

The ambiguity of the relationship between protein stability and function is exacerbated by the biases of the various stability prediction methods, which arise in their training, like overrepresentation of destabilising variants, dependence on crystal resolution and residue replacement asymmetry. Having observed protein-specific performance heterogeneity, we suggest that in the future focus could be shifted to identifying functional and structural properties of proteins, which could be most amenable to structure and stability-based prediction of mutation effects. Additionally, a recent work has showcased the use of homology models in structural analysis of missense mutation effects associated with disease, demonstrating utility that rivals experimentally derived structures, and thus expanding the possible resource pool that could be taken advantage of for structure-based disease prediction methods^28^. Further, our disease-associated mutations set likely contains variants causing disease through other mechanisms, that do not manifest through strong perturbation of the structure, making accurate evaluation impossible. To allow better stability-based predictors, it is important to have robust annotation of putative variant mechanisms, which is currently lacking due to non-existent experimental characterisation. We hope our results encourage new hybrid approaches, which make full use of the best available tools and resources to increase our ability to accurately prioritise putative disease mutations for further study, and elucidate the relationship between disease and stability changes.

## Methods

Pathogenic and likely pathogenic missense mutations were downloaded from the ClinVar^2^ database on 2019-04-17, while putatively benign variants were taken from gnomAD v2.1^1^. Any ClinVar mutations were excluded from the gnomAD set. We searched for human protein-coding genes with at least 10 ClinVar mutations occurring at residues present in a single high-resolution (< 2 Å) crystal structure of a protein that is monomeric in its first biological assembly in the Protein Data Bank. We excluded non-monomeric structures due to the fact that several of the computational predictors can only take a single polypeptide chain into consideration. All mutations and corresponding structures and predictions are provided in Table S1.

FoldX 5.0^71^ was run locally using default settings. Importantly, the ‘BuildModel’ function was first used to repair all structures. Ten replicates were performed for each mutation to calculate the mean.

The Rosetta suite (2019.14.60699 release build) was tested on structures first pre-minimised using the minimize_with_cst application and the following flags: -in:file:fullatom; -ignore_unrecognized_res - fa_max_dis 9.0; -ddg::harmonic_ca_tether 0.5; -ddg::constraint_weight 1.0; -ddg::sc_min_only false. The ddg_monomer application was run according to a rigid backbone protocol with the following argument flags: -in:file:fullatom; -ddg:weight_file ref2015_soft; -ddg::iterations 50; -ddg::local_opt_only false; -ddg::min_cst false; -ddg::min true; -ddg::ramp_repulsive true;-ignore_unrecognized_res.

Predictions by ENCoM, DUET and SDM were extracted from the DynaMut results page, as it runs them as parts of its own scoring protocol. mCSM values from DynaMut coincided perfectly with values from the separate mCSM web server, and thus the server values were used, as DynaMut calculations yielded less results due to failing on more proteins.

All other stability predictors were accessed through their online webservers with default settings by employing the Python RoboBrowser web scrapping library. Variant effect predictors were run in the same way as described in our recent benchmarking study^10^.

Method performance was analysed in R using the PRROC and pROC packages, and AUC curve differences were statistically assessed through 10,000 bootstraps using the roc.test function of pROC. For DynaMut, I-Mutant 3.0, mCSM, SDM, SDM2 and DUET, the sign of the predicted stability score was inverted to match the convention of increased stability being denoted by a negative change in energy.

## Acknowledgements

J.A.M. is supported by an MRC Career Development Award (MR/M02122X/1) and is a Lister Institute Research Prize Fellow. We thank Benjamin Livesey for his help with running the variant effect predictors.

## References

1. Karczewski, K. J. et al. The mutational constraint spectrum quantified from variation in 141,456 humans. Nature 581, 434–443 (2020).

2. Landrum, M. J. et al. ClinVar: Public archive of relationships among sequence variation and human phenotype. Nucleic Acids Res. 42, 980–985 (2014).

3. Gulilat, M. et al. Targeted next generation sequencing as a tool for precision medicine. BMC Med. Genomics 12, 1–17 (2019).

4. Suwinski, P. et al. Advancing personalized medicine through the application of whole exome sequencing and big data analytics. Front. Genet. 10, 1–16 (2019).

5. Katsonis, P. et al. Single nucleotide variations: Biological impact and theoretical interpretation. 23, 1650–1666 (2014).

6. Stenson, P. D. et al. The Human Gene Mutation Database: towards a comprehensive repository of inherited mutation data for medical research, genetic diagnosis and next-generation sequencing studies. Hum. Genet. 136, 665–677 (2017).

7. Niroula, A. & Vihinen, M. Variation Interpretation Predictors: Principles, Types, Performance, and Choice. Hum. Mutat. 37, 579–597 (2016).

8. Thusberg, J., Olatubosun, A. & Vihinen, M. Performance of mutation pathogenicity prediction methods on missense variants. Hum. Mutat. 32, 358–368 (2011).

9. Kato, S. et al. Understanding the function–structure and function–mutation relationships of p53 tumor suppressor protein by high-resolution missense mutation analysis. Proc. Natl. Acad. Sci. 100, 8424–8429 (2003).

10. Livesey, B. J. & Marsh, J. A. Using deep mutational scanning to benchmark variant effect predictors and identify disease mutations. Mol. Syst. Biol. (2020) doi:10.15252/msb.20199380.

11. Richards, S. et al. Standards and Guidelines for the Interpretation of Sequence Variants: A Joint Consensus Recommendation of the American College of Medical Genetics and Genomics and the Association for Molecular Pathology. Genet. Med. 17, 405–424 (2015).

12. Khan, S. & Vihinen, M. Performance of protein stability predictors. Hum. Mutat. 31, 675–684 (2010).

13. Potapov, V., Cohen, M. & Schreiber, G. Assessing computational methods for predicting protein stability upon mutation: Good on average but not in the details. Protein Eng. Des. Sel. 22, 553–560 (2009).

14. Pucci, F., Bernaerts, K. V., Kwasigroch, J. M. & Rooman, M. Quantification of biases in predictions of protein stability changes upon mutations. Bioinforma. Oxf. Engl. 34, 3659–3665 (2018).

15. König, E., Rainer, J. & Domingues, F. S. Computational assessment of feature combinations for pathogenic variant prediction. Mol. Genet. Genomic Med. 4, 431–446 (2016).

16. Montanucci, L., Capriotti, E., Frank, Y., Ben-Tal, N. & Fariselli, P. DDGun: An untrained method for the prediction of protein stability changes upon single and multiple point variations. BMC Bioinformatics 20, 1–10 (2019).

17. Usmanova, D. R. et al. Self-consistency test reveals systematic bias in programs for prediction change of stability upon mutation. Bioinformatics 34, 3653–3658 (2018).

18. Lonquety, M. Benchmarking stability tools: comparison of softwares devoted to protein stability changes induced by point mutations prediction. Comput Sys Bioinf … 1–5 (2007).

19. Huang, P. S., Boyken, S. E. & Baker, D. The coming of age of de novo protein design. Nature 537, 320–327 (2016).

20. Marcos, E. & Silva, D. A. Essentials of de novo protein design: Methods and applications. Wiley Interdiscip. Rev. Comput. Mol. Sci. 8, 1–19 (2018).

21. Buß, O., Rudat, J. & Ochsenreither, K. FoldX as Protein Engineering Tool: Better Than Random Based Approaches? Comput. Struct. Biotechnol. J. 16, 25–33 (2018).

22. Nemethova, M. et al. Twelve novel HGD gene variants identified in 99 alkaptonuria patients: focus on ‘black bone disease’ in Italy. Eur. J. Hum. Genet. 24, 66–72 (2016).

23. Stanton, C. M. et al. Novel pathogenic mutations in C1QTNF5 support a dominant negative disease mechanism in late-onset retinal degeneration. Sci Rep 7, 12147 (2017).

24. Heyn, P. et al. Gain-of-function DNMT3A mutations cause microcephalic dwarfism and hypermethylation of Polycomb-regulated regions. Nat Genet 51, 96–105 (2019).

25. Holt, R. J. et al. De Novo Missense Variants in FBXW11 Cause Diverse Developmental Phenotypes Including Brain, Eye, and Digit Anomalies. Am. J. Hum. Genet. 105, 640–657 (2019).

26. Bhattacharya, R., Rose, P. W., Burley, S. K. & Prlić, A. Impact of genetic variation on three dimensional structure and function of proteins. PLoS ONE 12, 1–22 (2017).

27. Al-Numair, N. S. & Martin, A. C. R. The SAAP pipeline and database: tools to analyze the impact and predict the pathogenicity of mutations. BMC Genomics 14 Suppl 3, (2013).

28. Ittisoponpisan, S. et al. Can Predicted Protein 3D Structures Provide Reliable Insights into whether Missense Variants Are Disease Associated? J. Mol. Biol. 431, 2197–2212 (2019).

29. Wang, Z. & Moult, J. SNPs, protein structure, and disease. Hum. Mutat. 17, 263–270 (2001).

30. Alibés, A. et al. Using protein design algorithms to understand the molecular basis of disease caused by protein-DNA interactions: the Pax6 example. Nucleic Acids Res 38, 7422–7431 (2010).

31. Caswell, R. C., Owens, M. M., Gunning, A. C., Ellard, S. & Wright, C. F. Using Structural Analysis In Silico to Assess the Impact of Missense Variants in MEN1. J. Endocr. Soc. 3, 2258–2275 (2019).

32. Abildgaard, A. B. et al. Computational and cellular studies reveal structural destabilization and degradation of MLH1 variants in Lynch syndrome. 28.

33. Seifi, M. & Walter, M. A. Accurate prediction of functional, structural, and stability changes in PITX2 mutations using in silico bioinformatics algorithms. PLoS ONE 13, 1–23 (2018).

34. Scheller, R. et al. Toward mechanistic models for genotype–phenotype correlations in phenylketonuria using protein stability calculations. Hum. Mutat. 40, 444–457 (2019).

35. Alford, R. F. et al. The Rosetta All-Atom Energy Function for Macromolecular Modeling and Design. J. Chem. Theory Comput. 13, 3031–3048 (2017).

36. Dehouck, Y., Kwasigroch, J. M., Gilis, D. & Rooman, M. PoPMuSiC 2.1: A web server for the estimation of protein stability changes upon mutation and sequence optimality. BMC Bioinformatics 12, 151 (2011).

37. Capriotti, E., Fariselli, P. & Casadio, R. I-Mutant2.0: Predicting stability changes upon mutation from the protein sequence or structure. Nucleic Acids Res. 33, 306–310 (2005).

38. Worth, C. L., Preissner, R. & Blundell, T. L. SDM - A server for predicting effects of mutations on protein stability and malfunction. Nucleic Acids Res. 39, 215–222 (2011).

39. Pandurangan, A. P., Ochoa-Montaño, B., Ascher, D. B. & Blundell, T. L. SDM: A server for predicting effects of mutations on protein stability. Nucleic Acids Res. 45, W229–W235 (2017).

40. Pires, D. E. V., Ascher, D. B. & Blundell, T. L. MCSM: Predicting the effects of mutations in proteins using graph-based signatures. Bioinformatics 30, 335–342 (2014).

41. Pires, D. E. V., Ascher, D. B. & Blundell, T. L. DUET: A server for predicting effects of mutations on protein stability using an integrated computational approach. Nucleic Acids Res. 42, 314–319 (2014).

42. Parthiban, V., Gromiha, M. M. & Schomburg, D. CUPSAT: Prediction of protein stability upon point mutations. Nucleic Acids Res. 34, 239–242 (2006).

43. Laimer, J., Hiebl-Flach, J., Lengauer, D. & Lackner, P. MAESTROweb: A web server for structurebased protein stability prediction. Bioinformatics 32, 1414–1416 (2016).

44. Frappier, V., Chartier, M. & Najmanovich, R. J. ENCoM server: Exploring protein conformational space and the effect of mutations on protein function and stability. Nucleic Acids Res. 43, W395–W400 (2015).

45. Rodrigues, C. H. M., Pires, D. E. V. & Ascher, D. B. DynaMut: Predicting the impact of mutations on protein conformation, flexibility and stability. Nucleic Acids Res. 46, W350–W355 (2018).

46. Stefl, S., Nishi, H., Petukh, M., Panchenko, A. R. & Alexov, E. Molecular Mechanisms of Disease-Causing Missense Mutations. J. Mol. Biol. 425, 3919–3936 (2013).

47. Nishi, H. et al. Cancer Missense Mutations Alter Binding Properties of Proteins and Their Interaction Networks. PLoS ONE 8, e66273 (2013).

48. Greiner, M., Pfeiffer, D. & Smith, R. D. Principles and practical application of the receiver-operating characteristic analysis for diagnostic tests. Prev. Vet. Med. 45, 23–41 (2000).

49. Bromberg, Y. & Rost, B. Correlating protein function and stability through the analysis of single amino acid substitutions. BMC Bioinformatics 10, S8 (2009).

50. Grimm, D. G. et al. The evaluation of tools used to predict the impact of missense variants is hindered by two types of circularity. Hum Mutat 36, 513–523 (2015).

51. Vaser, R., Adusumalli, S., Leng, S. N., Sikic, M. & Ng, P. C. SIFT missense predictions for genomes. Nat. Protoc. 11, 1–9 (2016).

52. Ng, P. C. & Henikoff, S. SIFT: predicting amino acid changes that affect protein function. Nucleic Acids Res. 31, 3812–3814 (2003).

53. Henikoff, S. & Henikoff, J. G. Amino acid substitution matrices from protein blocks. Proc Natl Acad Sci U A 89, 10915–10919 (1992).

54. Schymkowitz, J. et al. The FoldX web server: An online force field. Nucleic Acids Res. 33, 382–388 (2005).

55. Guerois, R., Nielsen, J. E. & Serrano, L. Predicting changes in the stability of proteins and protein complexes: A study of more than 1000 mutations. J. Mol. Biol. 320, 369–387 (2002).

56. Witham, S., Takano, K., Schwartz, C. & Alexov, E. A missense mutation in CLIC2 associated with intellectual disability is predicted by in silico modeling to affect protein stability and dynamics. Proteins Struct. Funct. Bioinforma. 79, 2444–2454 (2011).

57. Takano, K. et al. An X-linked channelopathy with cardiomegaly due to a CLIC2 mutation enhancing ryanodine receptor channel activity. Hum. Mol. Genet. 21, 4497–4507 (2012).

58. Tokuriki, N., Stricher, F., Serrano, L. & Tawfik, D. S. How protein stability and new functions trade off. PLoS Comput. Biol. 4, 35–37 (2008).

59. Frappier, V. & Najmanovich, R. J. A Coarse-Grained Elastic Network Atom Contact Model and Its Use in the Simulation of Protein Dynamics and the Prediction of the Effect of Mutations. PLoS Comput. Biol. 10, (2014).

60. Nisthal, A., Wang, C. Y., Ary, M. L. & Mayo, S. L. Protein stability engineering insights revealed by domain-wide comprehensive mutagenesis. Proc. Natl. Acad. Sci. 116, 16367–16377 (2019).

61. Montanucci, L., Martelli, P. L., Ben-Tal, N. & Fariselli, P. A natural upper bound to the accuracy of predicting protein stability changes upon mutations. Bioinformatics 35, 1513–1517 (2019).

62. David, A., Razali, R., Wass, M. N. & Sternberg, M. J. E. Protein-protein interaction sites are hot spots for disease-associated nonsynonymous SNPs. Hum. Mutat. 33, 359–363 (2012).

63. Bergendahl, L. T. et al. The role of protein complexes in human genetic disease. Protein Sci. 28, 1400–1411 (2019).

64. McEntagart, M. et al. A Restricted Repertoire of De Novo Mutations in ITPR1 Cause Gillespie Syndrome with Evidence for Dominant-Negative Effect. Am. J. Hum. Genet. 98, 981–992 (2016).

65. Williamson, K. A. et al. Recurrent heterozygous PAX6 missense variants cause severe bilateral microphthalmia via predictable effects on DNA– protein interaction. Genet. Med. (2019) doi:10.1038/s41436-019-0685-9.

66. Olijnik, A. et al. Genetic and functional insights into CDA-I prevalence and pathogenesis. Under revision at J Med Genet (2020).

67. Stein, A., Fowler, D. M., Hartmann-Petersen, R. & Lindorff-Larsen, K. Biophysical and Mechanistic Models for Disease-Causing Protein Variants. Trends Biochem. Sci. 44, 575–588 (2019).

68. Baugh, E. H. et al. Robust classification of protein variation using structural modelling and large-scale data integration. Nucleic Acids Res. 44, 2501–2513 (2016).

69. Ancien, F., Pucci, F., Godfroid, M. & Rooman, M. Prediction and interpretation of deleterious coding variants in terms of protein structural stability. Sci. Rep. 8, 1–11 (2018).

70. Kellogg, E. H., Leaver-Fay, A. & Baker, D. Role of conformational sampling in computing mutation-induced changes in protein structure and stability. Proteins Struct. Funct. Bioinforma. 79, 830–838 (2011).

71. Delgado, J., Radusky, L. G., Cianferoni, D. & Serrano, L. FoldX 5.0: working with RNA, small molecules and a new graphical interface. Bioinformatics 35, 4168–4169 (2019).

